# Phytohemagglutinin-Activated CAR-T Cells: Prolonged Persistence and Enhanced Anti-Tumor Response in CD19-Specific Acute Lymphoblastic Leukemia

**DOI:** 10.1101/2023.07.15.547808

**Authors:** Berranur Sert, Gamze Gulden, Tarik Teymur, Yasin Ay, Raife Dilek Turan, Onur Mert Unaldi, Elanur Güzenge, Hamza Emir Erdil, Sevim Isik, Pınar Oz, Ilknur Bozkurt, Tahire Arpacı, Osman Kamalı, Ercüment Ovalı, Nevzat Tarhan, Cihan Tastan

## Abstract

In recent times, chimeric antigen receptor (CAR)-T cell therapy has shown rapid advancements and gained clinical approval for use in cancer immunotherapy. CAR, a synthetic receptor integrated into autologous T cells, has yielded highly successful results in patients with leukemia. The significant potential of CAR-T cells has been validated through clinical trials in adult and pediatric cancer treatments. Our therapy developed specifically for CD19-specific Acute Lymphoblastic Leukemia (ALL) has shown promising results in in vitro and in vivo tests. To enhance the response against cancer, provide a bistimulatory effect, and increase stability, we designed two different CAR structures specific to CD19. These designs incorporate the CD28 and 41BB costimulatory domains. Through in vitro analysis, we evaluated the population ratios and cytotoxic activities of Central Memory T cells (TCM) and Stem Cell Memory T cells (TSCM) in CAR-T (CAR1928-T and CAR19BB-T) cells. Our initial design, CAR1928-T, produced an effective anti-tumor response. With our second design, CAR19BB-T, we not only achieved an anti-tumor effect but also conferred memory capabilities, leading to a comprehensive treatment approach. We demonstrated that CAR-T cells produced using Phytohemagglutinin (PHA) exhibited increased persistence in vitro and in vivo compared to anti-CD3 and anti-CD28 stimulation. The use of PHA to activate CAR19BB-T cells developed a long-lasting and effective CAR-T cell production method in vivo using cancerous animal models. CAR-T cell-treated mice survived tumor-free for up to 60 days, surpassing the survival of mice that received tumors only. Additionally, CAR19BB-T cell production with PHA remained stable over time. These results highlight a novel CAR-T cell production approach with a high-memory T cell profile capable of delaying or preventing cancer relapse. We optimized the method for the production of long-term and effective CAR-T cells and tested it in preclinical experiments. As a result, it was demonstrated that CAR-T cells generated with PHA, when administered as a co-stimulatory dose, can provide continuous proliferation and long-term persistence without compromising their anti-cancer efficacy. Preclinical studies have been completed to obtain valuable data for enhancing the long-term effectiveness of CAR-T therapy in clinical trials and transitioning to clinical applications.

## INTRODUCTION

According to data released by the World Health Organization (WHO), approximately 18-20 million people were diagnosed with cancer in 2018, and 10 million people lost their lives to cancer (World Health Organization, 2018). The most important strategies for cancer control are prevention and early diagnosis, along with the development of new treatment methods. However, due to the limited effectiveness of T-cell-based therapeutic approaches, such as immunotherapy, which utilize and enhance the patient’s immune system, research continues to address the heterogeneity of cancer cells (American Cancer Society, 2021; Mellman et al., 2011). Immature T lymphocytes are characterized by high expression of the transmembrane phosphatase CD45RA isoform (Janeway et al., 2001). The TN lymphocyte subset lacks the expression of CD45RO, CD95, CD11a, CD122, CD31, and KLRG1, and it is also characterized by high proliferative capacity (Cantillo et al., 2022; Sallusto et al., 1999).

T cells exhibit a natural immune response against tumors and demonstrate anti-tumor activity. T cells utilize a range of mechanisms to recognize and eliminate tumor cells (Vesely et al., 2011). The anti-tumor activity of T cells occurs through multiple mechanisms during their interactions with tumor cells (Vesely et al., 2011). Co-stimulatory molecules are proteins found on the surface of antigen-presenting cells that enhance T-cell activation and contribute to the initiation of an appropriate immune response (Chen & Flies, 2013). CD28: CD28 is the best-known co-stimulatory molecule on T cells, and it enhances T cell activation by interacting with the B7 molecules provided by antigen-presenting cells (Croft, 2003). 4-1BB: By binding to its ligand, 4-1BBL, 4-1BB enhances T cell activation and augments T cell function (Vinay & Kwon, 2012). The 4-1BB molecule also promotes the generation of long-lived memory T cells (Vinay et al., 2012). Therefore, the 4-1BB co-stimulatory receptor can be used in immunotherapy strategies to enhance anti-tumor responses (Zhang et al., 2016; Curran et al., 2010). Consequently, CAR-T cells incorporating both CD28 and 4-1BB domains demonstrate anti-tumor activity. The selection of domains should be based on the specific tumor type, patient characteristics, and treatment goals. Future clinical studies and more data will provide a clearer understanding of domain selection.

Anti-CD3&anti-CD28 is designed to activate and expand human T cells from enriched T cell populations or peripheral blood mononuclear cells (PBMCs) by providing signals through both the CD3 and CD28 pathways (Kinter et al., 2008). PHA is a lectin derived from red kidney beans that bind to T cell membranes and stimulates metabolic activity and cell division (Movafagh et al., 2011). PHA is a common polyclonal stimulator used in retroviral transduction protocols (Duarte et al., 2002). The concept of chimeric antigen receptor (CAR) engineering has been in use for over 25 years, and its effectiveness is still being investigated (Zhang et al., 2017; Gross Waks, 2018). Chimeric antigen receptor T (CAR-T) cell therapy is a customizable cell therapy approach used in immunotherapy (Zhang et al., 2017; Wang et al., 2018). Since 2017, the U.S. Food and Drug Administration (FDA) has approved CAR-T cell therapies for CD19+ hematological cancers (FDA, 2017; Maude et al., 2018). Applications of CAR-T cells continue to progress in the clinical setting, particularly in the treatment of hematological malignancies (Patterson et al., 2020).

Literature studies have shown that high proportions of central memory T cells (TCM) and stem cell-like memory T cells (TSCM) within CAR-T cell populations are associated with the necessary prerequisites for the effectiveness of immunotherapy, including continuous proliferation and long-term persistence (Blaeschke et al., 2016; Blaeschke et al., 2018). Therefore, the goal has been to increase the TCM and TSCM populations expressing CAR in order to achieve long-term and cytotoxic (anti-tumor) efficacy of CAR-T cells in vitro culture or in vivo cancer animal models. To increase the TCM and TSCM sub-populations within CAR-T cells, modifications have been made to the CAR gene construct (using 4-1BB or CD28-based CAR) and/or cell activation methods (Phytohemagglutinin/anti-CD3&anti-CD28) to develop a long-term, stable, and cytotoxic effective CAR-T cell production method. Studies have shown that CAR19BB-T cells have longer expression and more effective effector function compared to CAR1928-T cells (Lee et al., 2015; Maude et al., 2014; Sadelain et al., 2013). In this study, CAR-T cells produced with two different CAR constructs (aCD19 scFv-CD8-(CD28 or 41BB)-CD3z-EGFRt/GFP) were immunophenotyped, and the proportions of TCM and TSCM populations, immunodeficiency phenotypes, and cytotoxic efficacy were compared. Secondly, Phytohemagglutinin (PHA), a lectin that binds to T cell membranes and enhances metabolic activity and cell division, was tested to increase the TCM population. Although literature studies have indicated that PHA significantly increases TCM cells (Duarte et al., 2002), it has not been tested in CAR-T cell production and efficacy studies until now. Our study demonstrated that the CAR-T cell population generated with PHA had a higher proportion of TCM compared to CAR-T cells expanded with anti-CD3&anti-CD28, and it exhibited similar cytotoxic (anti-tumor) efficacy. Therefore, for the first time in the literature, the stability and efficacy of CAR-T cells produced with PHA were compared to CAR-T cells expanded with anti-CD3&anti-CD28 in an in vivo animal model.

This study aimed to culture CAR-T cells with PHA to confer a TCM and TSCM profile to these cells, enabling them to maintain long-term cytotoxic efficacy in both *in vitro* and *in vivo* cancer models. This paper optimized the method for the production of long-term and effective CAR-T cells and tested it in preclinical experiments. As a result, it was demonstrated that CAR-T cells generated with PHA, when administered as a co-stimulatory dose, can provide continuous proliferation and long-term persistence without compromising their anti-cancer efficacy. Preclinical studies have been completed to obtain valuable data for enhancing the long-term effectiveness of CAR-T therapy in clinical trials and transitioning to clinical applications.

## MATERIALS & METHODS

The CAR gene designs used in our study (aCD19 scFv-CD8-(CD28 or 4-1BB)-CD3z-EGFRt) consist of the sequence of an anti-CD19 monoclonal antibody’s (FMC63 clone) scFv (single-chain variable fragment) region, CD8 hinge region sequence, CD28 or 4-1BB transmembrane and co-stimulatory signaling domain sequences, and CD3z pro-activator intracellular domain sequence. The lentiviral vectors encoding CAR1928 (CD19-CD8-CD28-CD3z) and CAR19BB (CD19-CD8-41BBCD3z) were designed by our team and synthesized by GenScript. The envelope pCMV-VSV-G plasmid (Addgene #8454) and psPAX2 plasmid (Addgene #12260) necessary for lentivirus production were obtained from Addgene. The plasmids, including the CAR-encoding plasmids for genetic modification and the plasmids used for lentivirus packaging proteins (VSVG and psPAX2), were integrated into E. coli DH5α (NEB C2987H) strain and amplified. Plasmid DNAs were isolated using the Zymopure Plasmid Maxiprep Kit (Zymopure, Cat: D4202). The DNA concentration was measured as ng/μL using a Microplate ELISA Reader (FLUOstar Omega), and the purity was evaluated by checking if the A260/A280 ratio was between 1.8 and 2.0. For the control of isolated plasmids, DNA samples were loaded onto a 1% agarose gel prepared in 1X TAE buffer using a BIO-RAD gel electrophoresis system and run at 90 V for 60 minutes. Plasmid DNA samples with <%1 bacterial DNA contamination were used for lentivirus production. The isolated envelope, packaging, and CAR plasmids were transfected into HEK293T host cells (in Gibco DMEM HG medium containing 10% FBS, 1% penicillin/streptomycin, and L-Glutamine) using Polyethylenimine (PolyScience 23966-100) or FuGENE (Promega, Madison, WI, USA, E2311) transfection reagent to produce lentivirus. The packaged recombinant lentiviruses were collected from the supernatant of HEK293T cell cultures 72 hours after transfection. The produced CAR lentiviruses were concentrated using Lenti-X Concentrator (Takara Bio, Shiga, Japan, 631232) to increase the virus concentration (20X-100X). To filter out host-cell protein (HCP) residuals, the produced lentiviruses were diafiltered using an Amicon filter (100 kDa), and the virus samples that met the criteria of HCP ratio <200 ng/ml and negative for Mycoplasma hominis/genitalium were stored at -80°C.

### 1. Lentivirus Titration (Infection Unit; IFU/ml)

Jurkat cells were suspended in 100 μL RPMI medium containing 10% FBS, 1% penicillin/streptomycin, 1% non-essential amino acids, 1% sodium pyruvate, and 1% vitamins at a concentration of 10,000 cells per well. The Jurkat cells were added to U-bottom 96-well plates from A to H, with each well containing 100 μL of the medium. The wells were adjusted to have a volume of 150 μL with 10 μL, 3 μL, 1 μL, 0.3 μL, 0.1 μL, 0.03 μL, and 0.01 μL of aCD19 CAR lentivirus solution, which was 100X concentrated. The cells were incubated for 72 hours. The expression of EGFRt was determined using an anti-EGFR-FITC antibody (R&D Systems, Minneapolis, MN, USA, FAB10951G) or GFP expression analyzed by Cytoflex Flow Cytometer (Beckman Coulter, Brea, CA, USA, B5-R3-V0). The lentivirus concentration was confirmed by p24 ELISA Test in Infectious Units per milliliter (IFU/ml) and calculated using the following protocol.

### 2. T Cell Transduction and CAR-T Culture Conditions

Our in vitro and in vivo studies were approved by the Acibadem University and Acibadem Healthcare Institutions Medical Research Ethics Committee (ATADEK-2019-17/31). Peripheral blood mononuclear cells (PBMCs) were isolated from healthy adult blood samples obtained at the Transgenic Cell Technologies and Epigenetic Application and Research Center (TRGENMER) at Uskudar University. Blood samples from three healthy donors were combined with Ficoll (Paque PREMIUM 1.073, 17-5446-52) for isolation. PBMCs were isolated using density gradient centrifugation. Isolation of CD3+ T cells from PBMCs for CD4+ and CD8+ T cell isolation was performed using anti-CD3 microbeads (Miltenyi Biotech, Bergisch Gladbach, Germany, 130-050-101). Initial T cell activation was performed using anti-CD3/anti-CD28 microbeads (T Cell TransAct, Miltenyi Biotech, 130-111-160) and 10 μg/mL Phytohemagglutinin-M (PHA-M) (Roche 11082132001, 20 mg). Lentiviral transduction was performed by inoculating cells with Vectofusin 1 (10 μg/mL) (Miltenyi Biotech, 130-111-163). Cells were cultured in T cell medium (50 IU/mL IL-2, 10 ng/mL IL-7, 5 ng/mL IL-15, 10% Fetal Bovine Serum, and 1% penicillin/streptomycin) for 14 days with 3 MOI lentivirus. CAR expression levels were determined by Cytoflex Flow Cytometer analysis, either using an anti-EGFR-FITC antibody or by monitoring GFP expression. On the 14th day, a second activation and proliferation assay was performed using PHA and anti-CD3/anti-CD28.

### 3. In Vivo Anti-Tumor Formation in ALL Mouse Model

In vivo, anti-tumor efficacy experiments were conducted in Nude/NSG mice using RAJI cells (Burkitt lymphoma) transduced with lentiviruses expressing mCherry and firefly luciferase (fLuciferase). The mCherry expression of RAJI cells was confirmed by flow cytometry (>97% mCherry positive). The in vivo cancer animal model consisted of 6 groups (**Figure 1**). To establish a tumor model, 500,000 fLuciferase-mCherry positive RAJI cells were injected into Nude/NSG mice. Tumor growth and development were monitored using bioluminescent imaging upon injection of D-Luciferin, Potassium Salt (Proven and Published™) 100 mg (Goldbio, LUCK-100, 1mg/mL). The bioluminescent signal emitted by the disseminated tumors was measured at 7-day intervals for 2 months using the IVIS Spectrum In Vivo Imaging System (PerkinElmer). The weight of the mice was recorded before each measurement.

**Figure 1.**
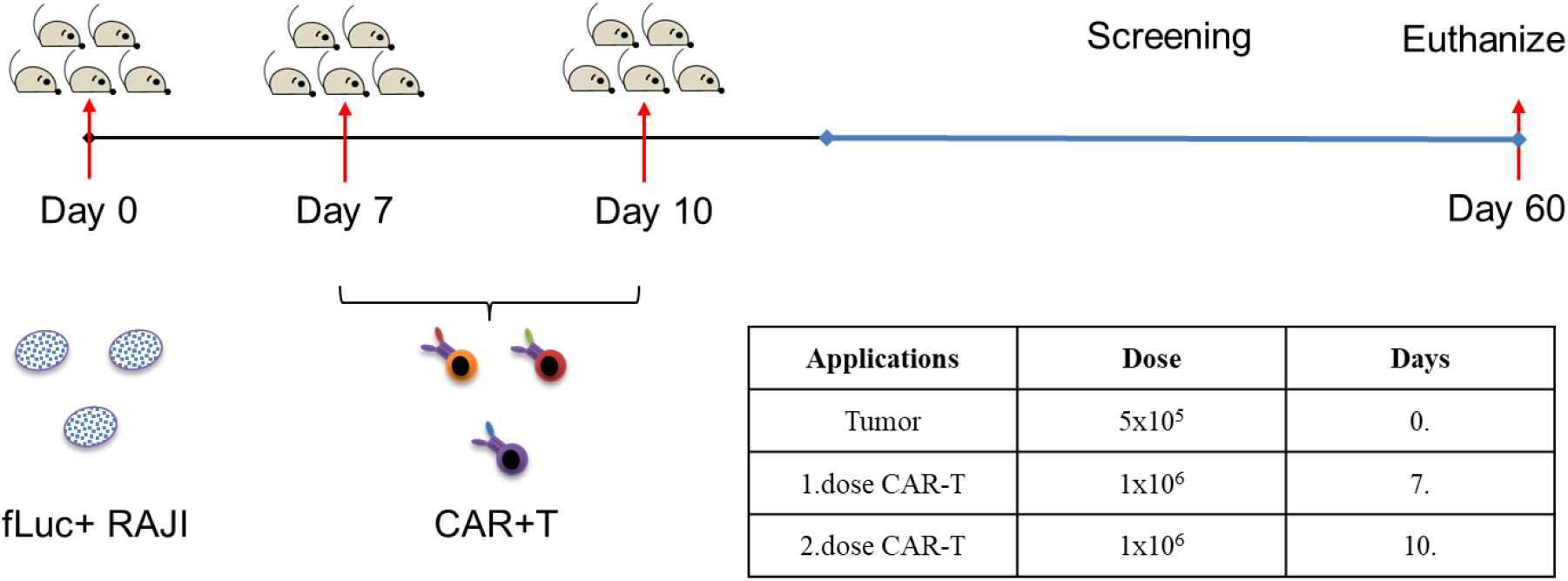
Injection administration schedule.

### 4. CAR-T Cell Application in the ALL Mouse Model

All animal studies have been approved by Acıbadem University Animal Experiments Local Ethics Committee (ACU-HADYEK; 11.13.2019). Tumor presence was observed on the seventh day based on mCherry expression, and CAR-T cell therapy was initiated on day 7. In the first group, only a tumor model was established, and on the 7th and 10th days, saline was injected intraperitoneally **(Figure 1)**.

In the treatment groups, there are three different CAR-T cell groups. In the second group, after tumor application, two doses of CAR19BB anti-CD3&anti-CD28 CAR-T (1×106 IP per mouse) were administered on the 7th and 10th days. In the third group, after tumor application, CAR19BB anti-CD3&anti-CD28 CAR-T (1×106 IP per mouse) was administered on the 7th day, and CAR19BB PHA (1×106 IP per mouse) was administered on the 10th day. In the fourth group, after tumor application, two doses of CAR1928 PHA CAR-T (1×106 IP per mouse) were administered on the 7th and 10th days. The fifth group did not receive any tumor applications. The study plan is presented in **Figure 2**. Tumor growth and development were monitored using bioluminescence imaging after D-Luciferin injection.

**Figure 2.**
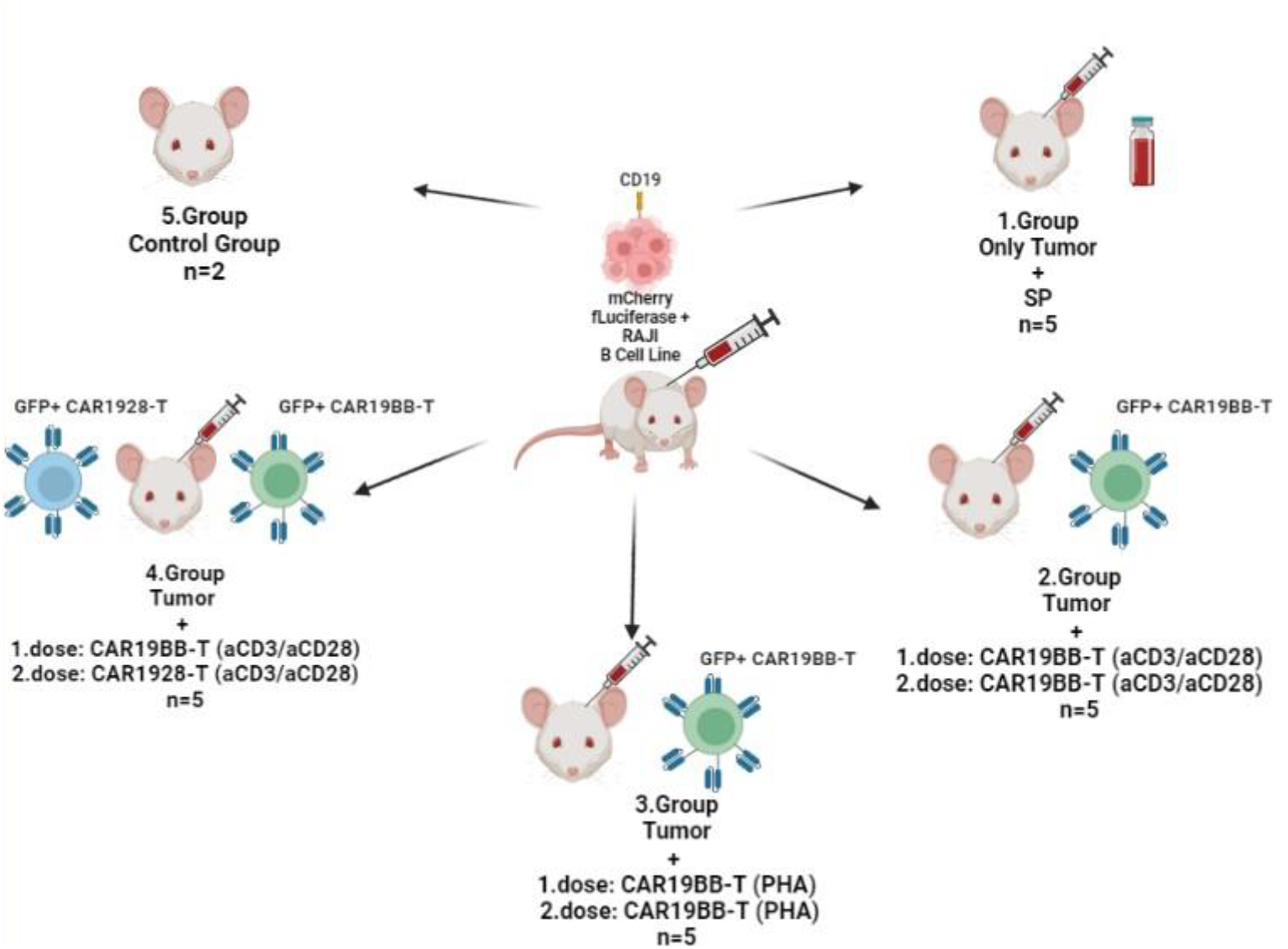
In vivo CAR-T cell Efficacy Studies in ALL Nude/NSG Mouse Model.

### 5. ALL Model Mice Blood Analyses

During the preclinical study, 1 cc blood samples were obtained intraventricularly from all mouse groups during sacrifice, and the following analyses were performed: complete blood count (hemogram), ALT, AST, BUN, creatinine, calcium, and albumin levels. The blood analyses were conducted by obtaining services from the Medical Biochemistry Laboratory of NP Üsküdar Brain Hospital. Additionally, genomic DNA isolation was performed from 0.05 cc blood samples using the protocol described below, and the presence of CAR-T cells was evaluated using HIV-1 PCR testing.

### 6. Genomic DNA Isolation

The required amount of Elution Buffer was incubated at 60°C throughout the steps. 400 µl of Lysis Solution SLS and 30 µl of Proteinase K were added to each 0.05 cc whole blood sample. The samples were vortexed for 10 seconds and incubated at 60°C for 10 minutes. Then, 700 µl of Binding Solution BL was added to the samples. The samples were centrifuged at 11,000g (∼12,000 rpm) for 1 minute using Receiver and Spin columns. The Spin Filter was opened, and 400 µl of Washing Solution C was added. The samples were centrifuged at 11,000g for 1 minute. Subsequently, 600 μl of Washing Solution BS was added. The samples were centrifuged at 11,000g for 1 minute. Then, 200 µl of Elution Buffer was added, and the samples were incubated at room temperature for 2 minutes. The samples were centrifuged at 11,000g for 1 minute. The isolated DNA was stored at -20°C.

### 7. HIV-1 Real-Time PCR Test

DIAGNOTECH HIV-1 Real-Time PCR is an in vitro nucleic acid amplification test used for the quantitative detection of HIV-1. The test kit contains specific primers and fluorescent probes targeting conserved regions of the HIV-1 genome, including the LTR sequence found in lentiviruses. These probes enable the precise detection of the integrated LTR sequence in the genome of clinical samples using real-time fluorescent PCR technique. For the purpose of assessing the survival capacity of CAR-T cells injected for therapeutic purposes in vivo animal models, blood samples were collected during sacrifice from mice treated with CAR-T cells, and genomic DNA isolation was performed. Subsequently, the isolated DNA was subjected to real-time PCR (RT-PCR) testing, a single-step procedure in which the amplified product is detected using fluorescent dyes. Monitoring the fluorescence intensities during RT-PCR allows the detection and quantification of the accumulated product. This monitoring is performed by binding fluorescent dyes specific to oligonucleotide probes that are specific to the amplified product. This test was used in this study to detect LTR sequences that are integrated into the genome of CAR-T cells. The DIAGNOTECH HIV-1 Real-Time PCR Kit (Catalog No: DB0107100) was performed according to the manufacturer’s instructions. The 20 µl Multiplex PCR Master Mix included in the kit was mixed with 20 µl Genomic DNA and dispensed into the strips.

### 8. ALL Mouse Model: Sacrifice and Tissue Sectioning

For all mouse groups, mice were anesthetized using isoflurane inhalation and sacrificed by cervical dislocation for preclinical blood analyses. Liver, kidney, spleen, lung, and brain tissues were collected from each mouse and fixed in 4% paraformaldehyde (Thermo Scientific, J19943-K2, Lot: 214182) for 3 days at +4°C. After fixation, the tissues were transferred to 10% sucrose solution (Multicell, Cat: 800-081-LG) and incubated for 24 hours at +4°C. Following incubation, the tissues were transferred to 20% sucrose solution and incubated for 48 hours at +4°C. After 48 hours, the tissues were transferred to a 30% sucrose solution for preservation. Before tissue sectioning, the tissues were frozen using a cryo-gel embedding compound (Surgipath Cryo-Gel Embedding Compound for Frozen Section Leica, Ref: 39475237). Tissue sections with a size of 15-20 microns were obtained using a cryostat device (Leica CM1950 Cryostat). The obtained tissue sections were placed onto positively charged slides (PATOLAB, 1650139) and fixed in 100% ethanol (TEKKIM, TK.200650.05001) for 10 minutes. After fixation, the slides were briefly dipped in ddH2O and stored in PBS solution (CAPRICORN SCI, PBS-1A) at +4°C.

### 9. Histochemical Staining in ALL Mouse Model

Hematoxylin and Eosin histochemical staining were performed on the collected tissue sections from all mouse groups to examine necrosis, tissue integrity, and structural changes in cellular elements. The slides were taken out of the PBS solution and incubated in ddH2O for 1 minute. After incubation, the slides were dipped in Hematoxylin (Merck, Cat: 105174) for 2-3 seconds, then placed back into ddH2O and incubated for 1 minute. Next, the slides were immersed in 70% ethanol (TEKKIM, TK.200650.05001) for 30 seconds. This was followed by immersion in 95% ethanol for 30 seconds. Subsequently, the slides were placed in Eosin (Merck, Cat: 1098441000) for 5 seconds. After incubation in Eosin, the slides were incubated in 95% ethanol for 15 seconds, followed by incubation in 100% ethanol for 15 seconds. After incubation, the samples were briefly immersed in xylene (TEKKIM, Cat: TK.090270.05003) for 3 seconds. Then, 1 drop of Entellan (Merck, Cat: 107961) was added to the tissues, covered with a cover glass lamella (Menzel, Cat: M-24601), and stored at +4°C for imaging.

### 10. Statistical Analysis

Single/two-tailed t-tests were performed using SPSS software. Outliers were not excluded from any of the statistical tests, and each data point represents an independent measurement. Bar graphs represent the mean and standard deviation or mean with standard deviation. The significance threshold for all tests was set at *p < 0.05. (Ns: non-significant)

## RESULTS

Evaluation of the in vivo stability and efficacy capacities of CAR-T cells developed under different production conditions can provide new approaches for CAR-T cell production and application. For this purpose, CAR19BB-T and CAR1928-T cells produced by activation with PHA or anti-CD3&anti-CD28 were aimed to be evaluated in an in vivo study using a CD19+ B-cell cancer model in mouse groups. To simulate the CD19+ ALL cancer model in the in vivo mouse groups, a >99% fLuciferase+ mCherry+ RAJI cell line was developed through lentiviral transduction. The development of the ALL cancer model in the RAJI-injected mouse groups was monitored over nine weeks using bioluminescence (**Figure 3A**) and mCherry fluorescence (**Figure 3B**) channels. After the detection of mCherry RAJI formation in the fluorescence channel on day 7, CAR19BB-T and CAR1928-T cells produced by activation with PHA or anti-CD3&anti-CD28 were injected on days 7 and 10 (**Figure 3B**). When evaluating cancer development in the "only tumor, CAR19BB-T (anti-CD3&anti-CD28), CAR19BB-T (PHA), and CAR1928-T (anti-CD3&anti-CD28)" groups in the bioluminescence and mCherry channels, cancer development continued until day 35 in the only tumor group, while in the group injected with CAR19BB-T (anti-CD3&anti-CD28), cancer development persisted for two months in one mouse (n=5) (**Figure 3A and 3B**). In the CAR19BB-T (PHA) and CAR1928-T (anti-CD3&anti-CD28) groups, RAJI cancer development was not observed in the bioluminescence channel and disappeared in the mCherry channel after CAR-T cell injections on day 7 (**Figure 3A and 3B**). However, starting from day 28, all mice with advanced cancer development were sacrificed in the only tumor group from day 35 onwards (**Figure 3A**). These results demonstrate that CAR-T cell injections suppressed CD19+ cancer development in all groups for two months in vivo tests. Additionally, CAR-T cells encoding GFP were followed in the GFP fluorescence channel for two months in the in vivo injection groups to assess the presence of CAR-T cells (**Figure 3C**). Following the injection day (day 7), CAR-T cell presence was similarly detected in all groups, especially in the CAR19BB-T (PHA) injected group, stability was observed to be maintained without a decrease after two months (**Figure 3C**). In the CAR19BB-T (anti-CD3&anti-CD28) and CAR1928-T (anti-CD3&anti-CD28) injected groups, an increase in CAR-T cells was observed after day 49, parallel to the increase in mCherry+ RAJI luminescence (**Figure 3C**). These results indicate that CAR19BB-T activated with PHA has higher stability compared to other groups.

**Figure 3.**
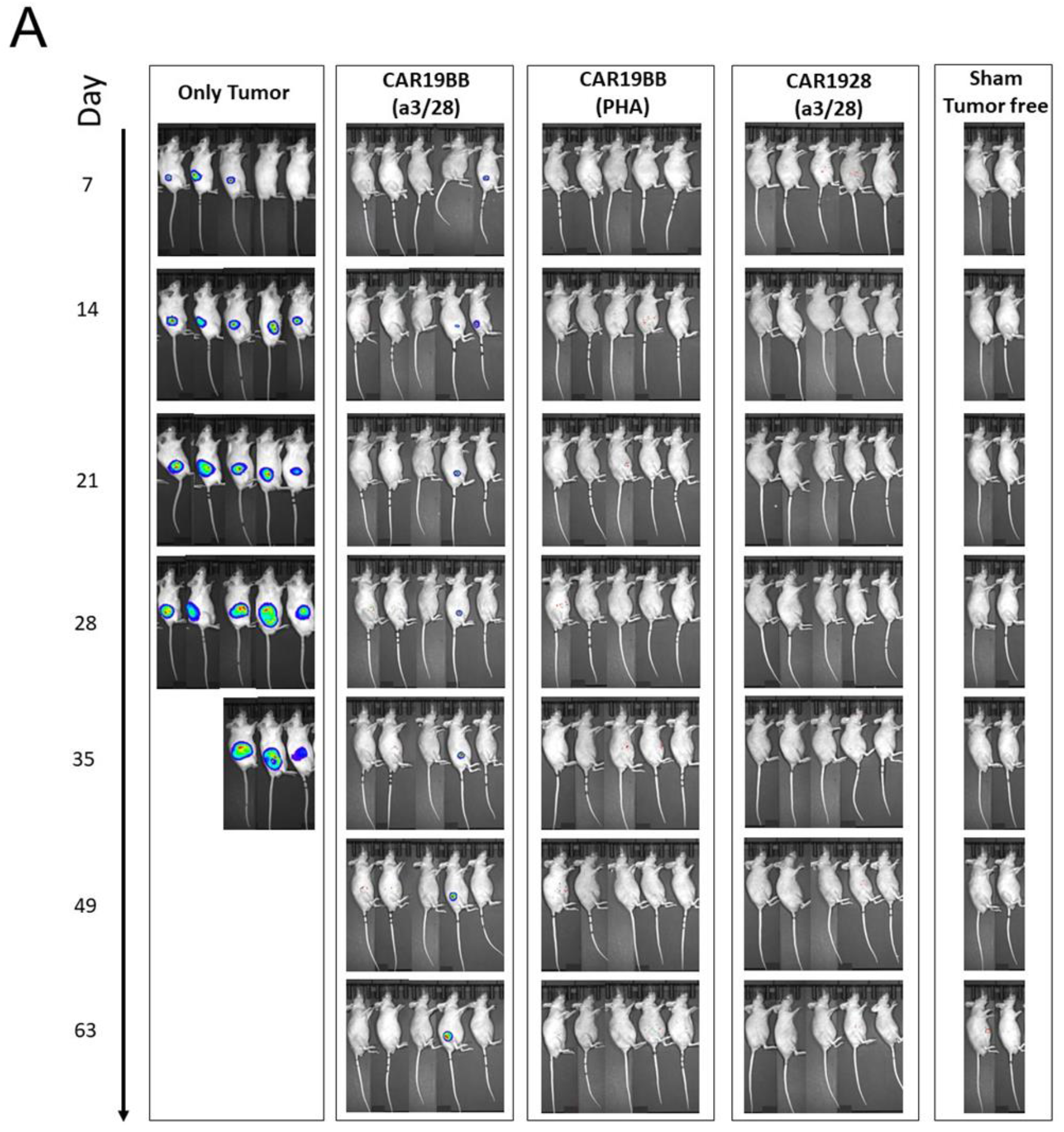
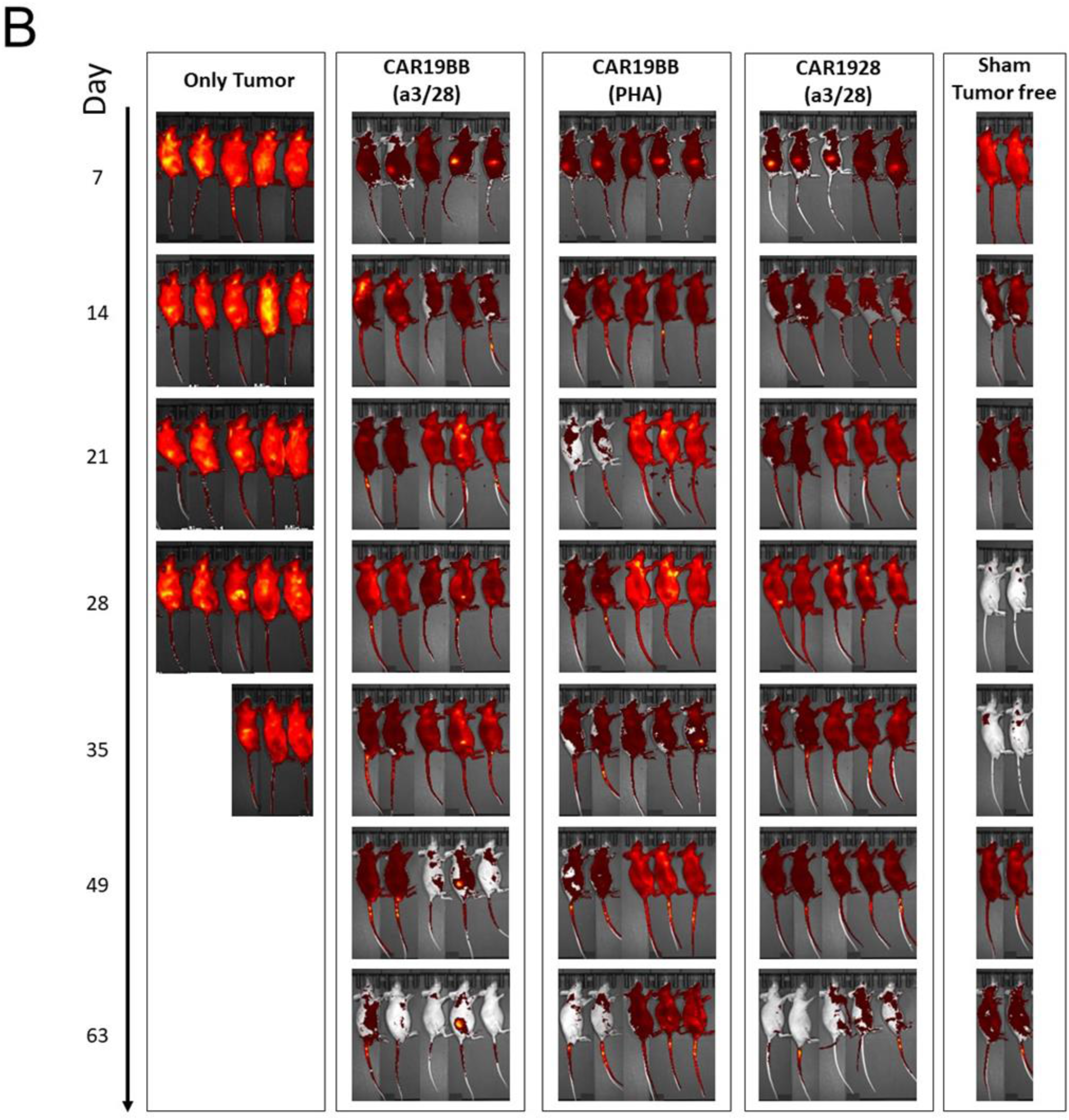
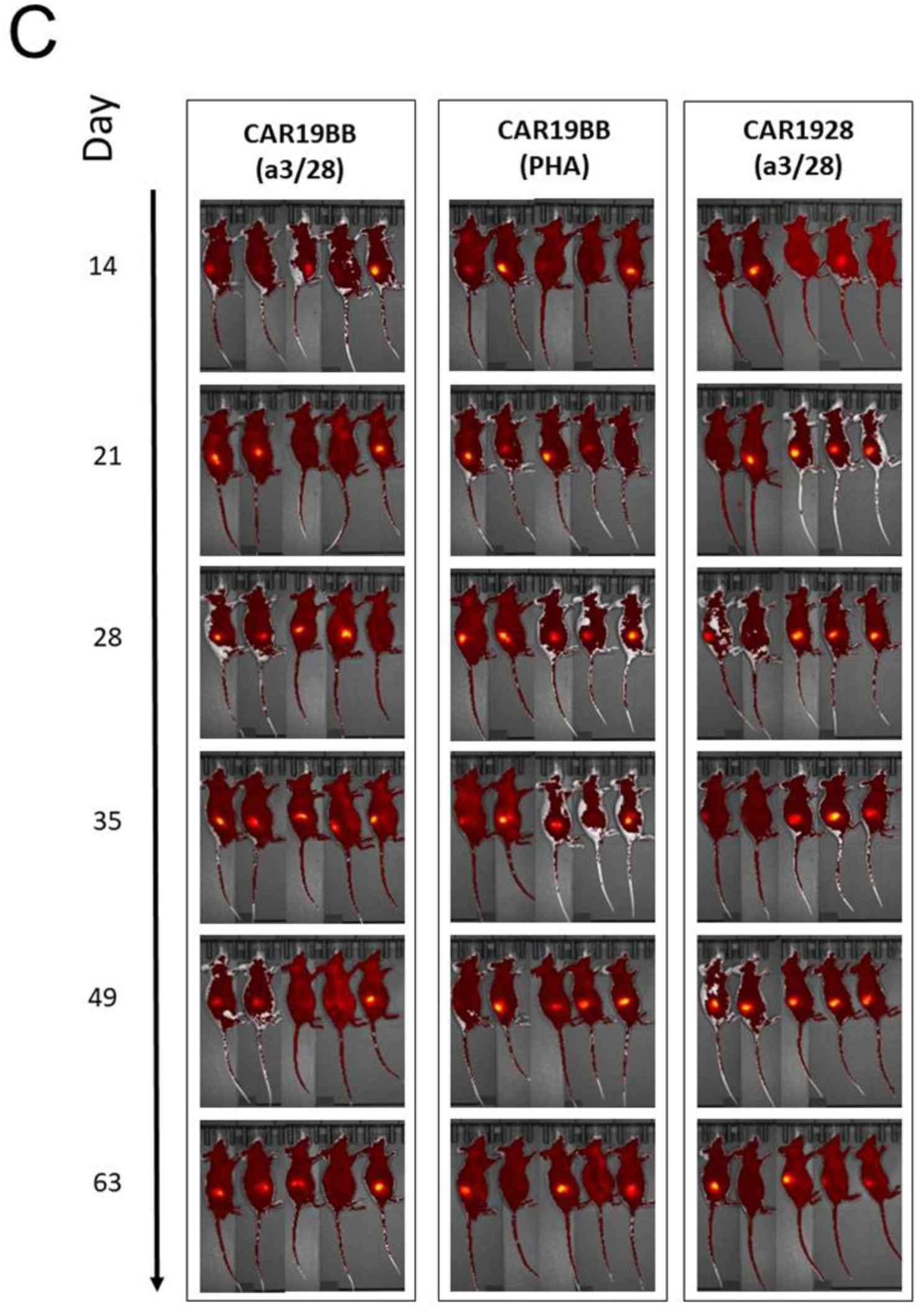
Time-dependent visualization of CAR19BB-T and CAR1928-T cell efficacy in an in vivo ALL cancer model in mice (7-63 days). (**A**) Bioluminescent imaging shows the luminescence of fLuciferase+ mCherry+ CD19+ RAJI cancer cells in the treatment groups following luciferin injection. (**B**) Fluorescent imaging shows the luminescence of fLuciferase+ mCherry+ CD19+ RAJI cancer cells in the treatment groups. (**C**) GFP fluorescent imaging shows the luminescence of GFP+ CAR-T cells in the treatment groups.

Imaging was initiated on day 14 after RAJI cell injection and the time-dependent quantification analysis of in vivo IVIS imaging was performed to investigate the monitoring of CD19+ RAJI and GFP+ CAR-T cells after in vivo injection for two months. It was observed that the luminescence of fLuciferase+ RAJI cells significantly increased, particularly from day 21 onwards until the sacrifice of the entire group on day 35, in the "only tumor" group compared to the CAR-T treatment groups (p=0.014) (**Figure 4A**). Additionally, it can be seen that the presence of CAR19BB-T (anti-CD3&anti-CD28) in the group with ongoing tumor development increased the average luminescence value within the group (**Figure 4A**). For the sensitive evaluation of RAJI cancer development, it is observed that the mCherry fluorescence luminescence assessments showed similarity to the fLuciferase bioluminescence luminescence, especially in the "only tumor" group. In all CAR-T treatment groups, the mCherry+ RAJI cell ratio significantly disappeared, especially from day 14 onwards (**Figure 4B**). Furthermore, the relapse of RAJI cancer cell line in the mCherry channel could also be observed in the fluorescence channel, starting from day 49 **(Figure 4B**). Another question we asked in the study is whether the stability of CAR-T cells activated with PHA can be increased compared to cells activated with anti-CD3&anti-CD28. In the GFP fluorescence luminescence analysis, it was observed that the CAR-T cell ratio in the group treated with CAR19BB-T (PHA) did not decrease throughout the 63 days (**Figure 4C**). In contrast, in the group where CAR19BB-T and CAR1928-T cells were administered with anti-CD3&anti-CD28, the CAR-T cell ratios significantly decreased compared to the CAR19BB-T (PHA) ratio, starting from day 28 (p=0.032) (**Figure 4C**). To evaluate the proliferation and stability capacity of CAR-T cells in the case of time-dependent ALL relapse, GFP CAR-T, and mCherry RAJI fluorescence luminescence analyses were evaluated together. In the group where CAR19BB-T (anti-CD3&anti-CD28) cell administration was performed, it was observed that RAJI cancer development was zeroed from day 14 onwards, and at the same time, the CAR19BB-T (anti-CD3&anti-CD28) cell ratio continued to decrease until day 49 (**Figure 4D**). Moreover, the increase in CD19+ mCherry+ RAJI cells from day 49 to 63 was parallel to the increase in the CAR19BB-T (anti-CD3&anti-CD28) cell ratio (**Figure 4D**). On the other hand, in the group where CAR19BB-T (PHA) was administered, while RAJI cancer development was zeroed on day 14, no decrease was observed in the CAR-T cell ratio until day 63 (**Figure 4E**). In the group where CAR1928-T (anti-CD3&anti-CD28) cell administration was performed, it was observed that RAJI cancer development was zeroed from day 21 onwards, and at the same time, the CAR1928-T (anti-CD3&anti-CD28) cell ratio continued to decrease until day 49 (**Figure 4F**). These results demonstrate that CAR19BB-T cells activated with PHA maintain their anti-cancer efficacy and have higher stability compared to CAR19BB-T or CAR1928-T cells produced by activation with anti-CD3&anti-CD28 over time.

**Figure 4.**
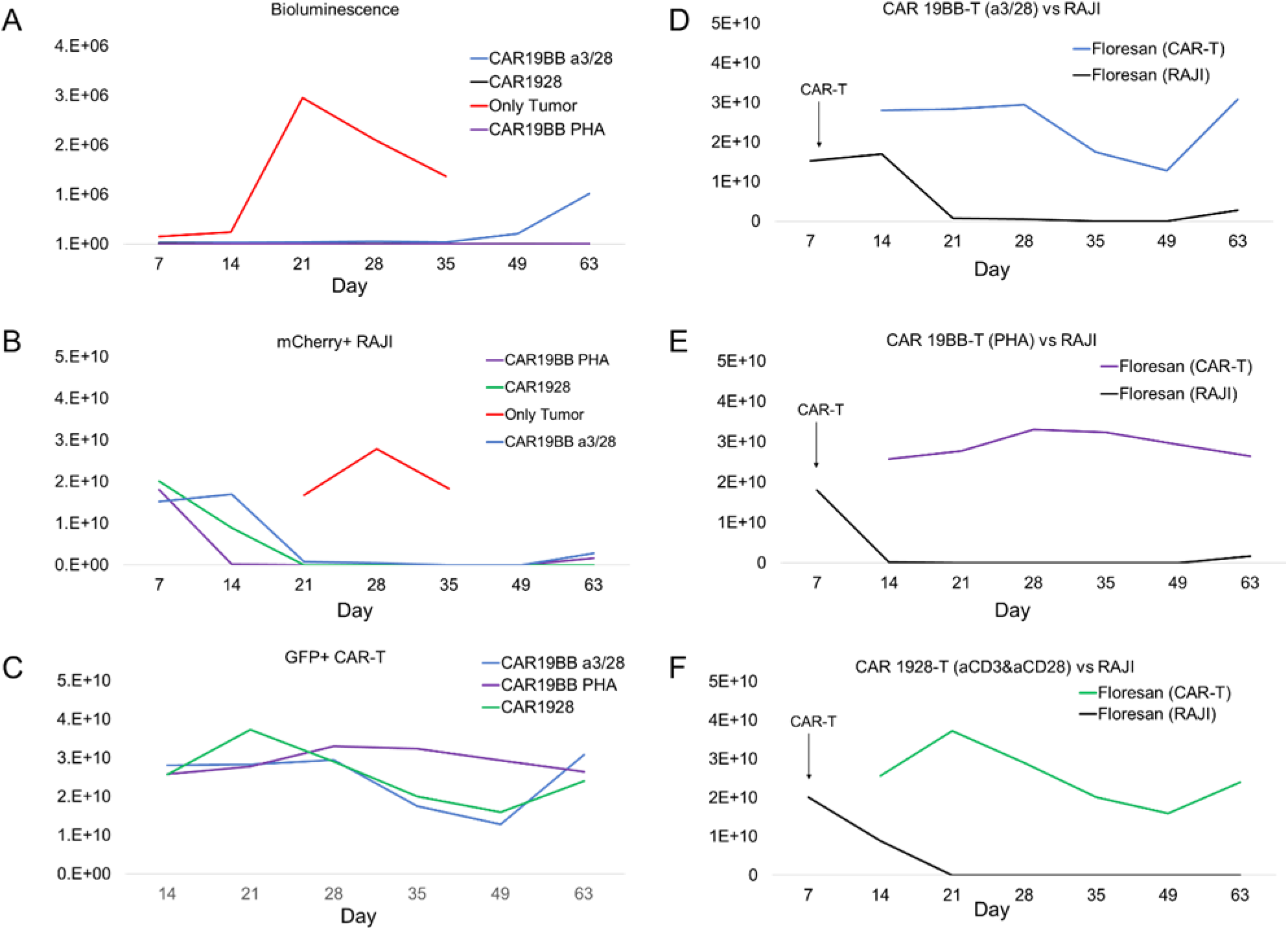
In vivo quantification of fLuciferase bioluminescence, mCherry, and GFP fluorescence. (**A**) The graph shows the time-dependent development (7-63 days) of fLuciferase+ RAJI cancer cells in the treatment groups in the bioluminescence luminescence channel in the ALL model mice. (**B**) The graph shows the time-dependent development (7-63 days) of mCherry+ RAJI cancer cells in the treatment groups in the mCherry fluorescence luminescence channel in the ALL model mice. (**C**) The graph shows the time-dependent presence (7-63 days) of GFP+ CAR-T cells in the treatment groups in the GFP fluorescence luminescence channel in the ALL model mice. The fluorescence luminescence channels demonstrate the comparison of mCherry+ RAJI cancer cell growth in the treatment groups with the presence of GFP+ CAR-T cells over time in (**D**) CAR19BB-T (anti-CD3&anti-CD28), (**E**) CAR19BB-T (PHA), and (**F**) CAR1928-T (anti-CD3&anti-CD28) groups.

A Kaplan-Meier curve was plotted for survival analysis in groups receiving CAR-T cell therapy. After in vivo study, all mice in the "only tumor" group died by day 35, while no decrease in overall survival was observed in the group receiving CAR-T cell therapy after a two-month analysis (**Figure 5A**). However, for all surviving mice, specific LTR-targeted RT-PCR analysis of CAR-T cells was performed using blood samples collected after sacrifice on day 63 to assess CAR-T cell stability. In **Figure 4C** and **4E**, the presence of CAR-19BB-T (PHA) was observed to be significantly higher compared to the groups receiving CAR19BB-T (p=0.007955) and CAR1928-T (p=0.036695) generated by activation with anti-CD3&anti-CD28, indicating statistical significance (**Figure 5B**). Based on these analyses, it has been demonstrated that CAR19BB-T cells activated with PHA maintain their anti-cancer efficacy and exhibit higher stability over time compared to other application approaches.

**Figure 5.**
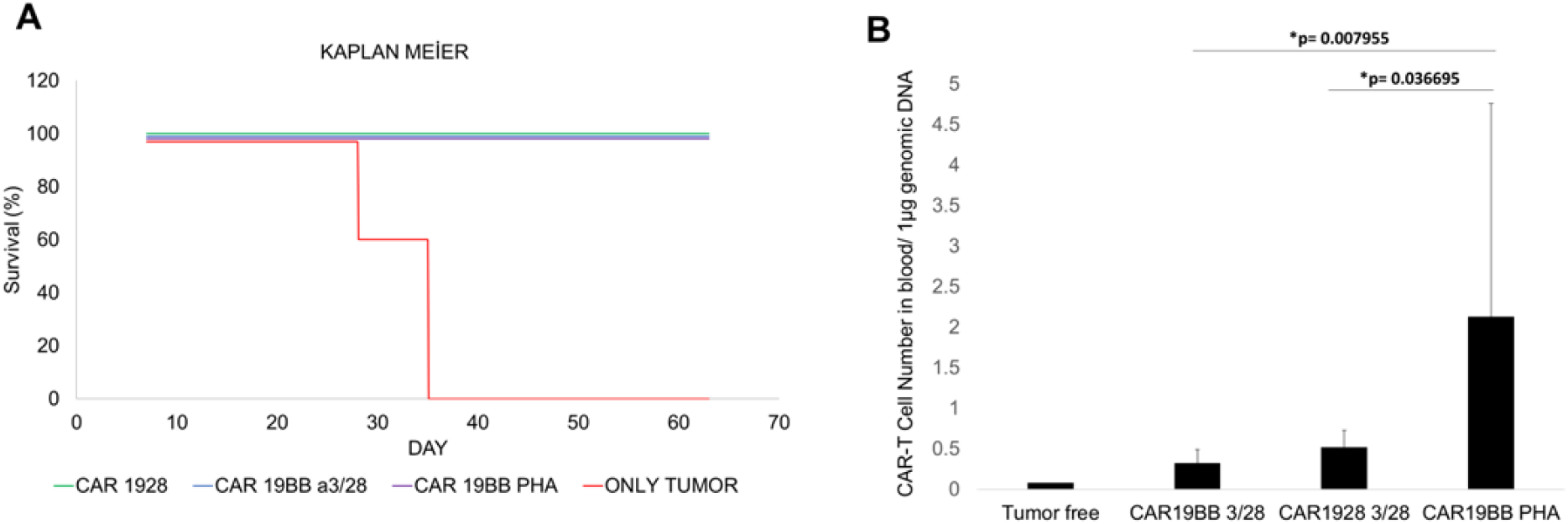
Kaplan-Meier Survival Analysis and In vivo PCR Detection of CAR-T Cells. (**A**) Survival analysis from day 7 to day 63 in mouse groups with ALL cancer models following CAR-T cell injections. (**B**) Bar graph represents the number of CAR-T cells in 1 μg of genomic DNA isolated after LTR-specific RT-PCR analysis of blood samples from ALL cancer mouse groups. Statistical significance is represented by one-sided t-test, * p < 0.05.

We evaluated liver, pancreas, and systemic inflammation in mouse groups where CAR-T cell therapy was terminated on the 63rd day, along with survival and anti-cancer analyses, by performing hemogram analysis and biochemical tests. Hemogram analysis showed no significant difference between the CAR-T treated groups and the tumor-free control group, and the values generally returned to physiological levels observed in the tumor-free control group compared to the only tumor group (**Figure 6A**). However, in the biochemical analysis, it was found that CAR-T treated groups had significantly decreased levels of Albumin and Creatinine, and increased levels of Urea compared to the only tumor groups (**Figure 6B and 6C**). To monitor liver function impairment, ALT and AST tests were conducted, and it was determined that CAR-T treated groups had a significant decrease compared to the only tumor groups and showed similar values to the tumor-free control group (**Figure 6D and 6E**). Finally, hemogram analysis revealed a significant decrease in platelet count similar to thrombocytopenia in the CAR-T treated groups (**Figure 6F**). Hemogram and biochemical analyses indicated that CAR19BB-T and CAR1928-T cells, produced by activation with PHA or anti-CD3/anti-CD28, similarly diverged from systemic physiology within the only tumor group, decreased inflammation, and approached the tumor-free control group.

**Figure 6.**
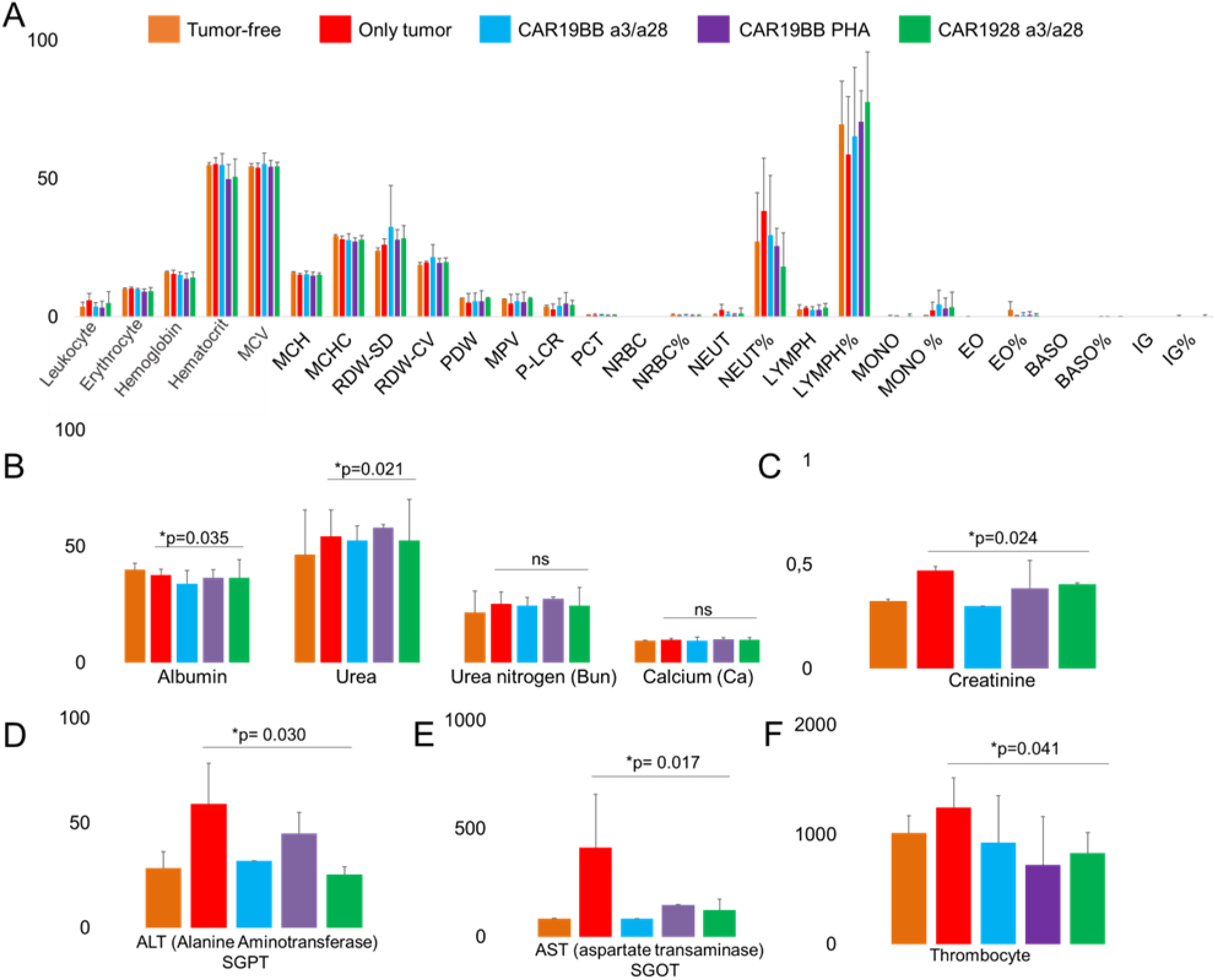
Hemogram and biochemical analysis in ALL cancer model mice after CAR-T cell therapy. (**A**) Bar graphs show the hemogram analysis on day 63 for the groups receiving CAR-T cell therapy and the tumor-free control group, as well as the hemogram analysis on day 35 for the only tumor group (n=5; group). The bar graphs depict the biochemical analysis on day 63 for the groups receiving CAR-T cell therapy and the tumor-free control group, as well as the biochemical analysis on day 35 for the only tumor group, including (**B**) Albumin (g/L), Urea (mg/dL), Urea Nitrogen (mg/dL), Calcium (mg/dL), (**C**) Creatinine (mg/dL), (**D**) ALT SGPT (U/L), and (**E**) AST SGOT (U/L). (**F**) Bar graphs show the Platelet count (103/uL) analysis on day 63 for the groups receiving CAR-T cell therapy and the tumor-free control group, as well as the Platelet count analysis on day 35 for the only tumor group (n=5; group). T-test statistical significance is represented as * p < 0.05.

Finally, after the sacrifice on day 63 of CAR-T cell-treated groups, tissue sections were obtained from the spleen, kidney, and liver for histopathological analysis, evaluating tissue integrity, necrosis, and inflammation. The obtained inverted microscope images following Hematoxylin & Eosin staining were compared with tissue sections from the only tumor group sacrificed on day 35. In spleen sections, it was observed that the number of white pulp (white arrow) areas, which are the sites of leukocyte production, was lower and the area was larger in the only tumor group, while in the CAR-T cell-treated groups, the number was higher and the area was narrower (**Figure 7A**). This result indicates that inflammation persists in the only tumor groups but decreases in the CAR-T cell groups. In the analysis of kidney sections, it was observed that the number of necrotic and injured areas (white arrow) was higher in the only tumor group compared to the CAR-T cell groups, indicating more pronounced cancer involvement (**Figure 7B**). Furthermore, when liver function impairment was investigated in pathology images, it was observed that structures resembling lymphocyte infiltration and necrotic formation (white arrow) were more pronounced in the only tumor group, suggesting a decrease in CAR-T cell-treated groups after biochemical analysis (**Figure 7C**). These results demonstrate that, regardless of the activation with PHA or anti-CD3/anti-CD28 and the structures of CAR1928 and CAR19BB, CAR-T cell-treated groups showed a reduction in necrosis and inflammation compared to the only tumor group, indicating improved tissue integrity.

**Figure 7.**
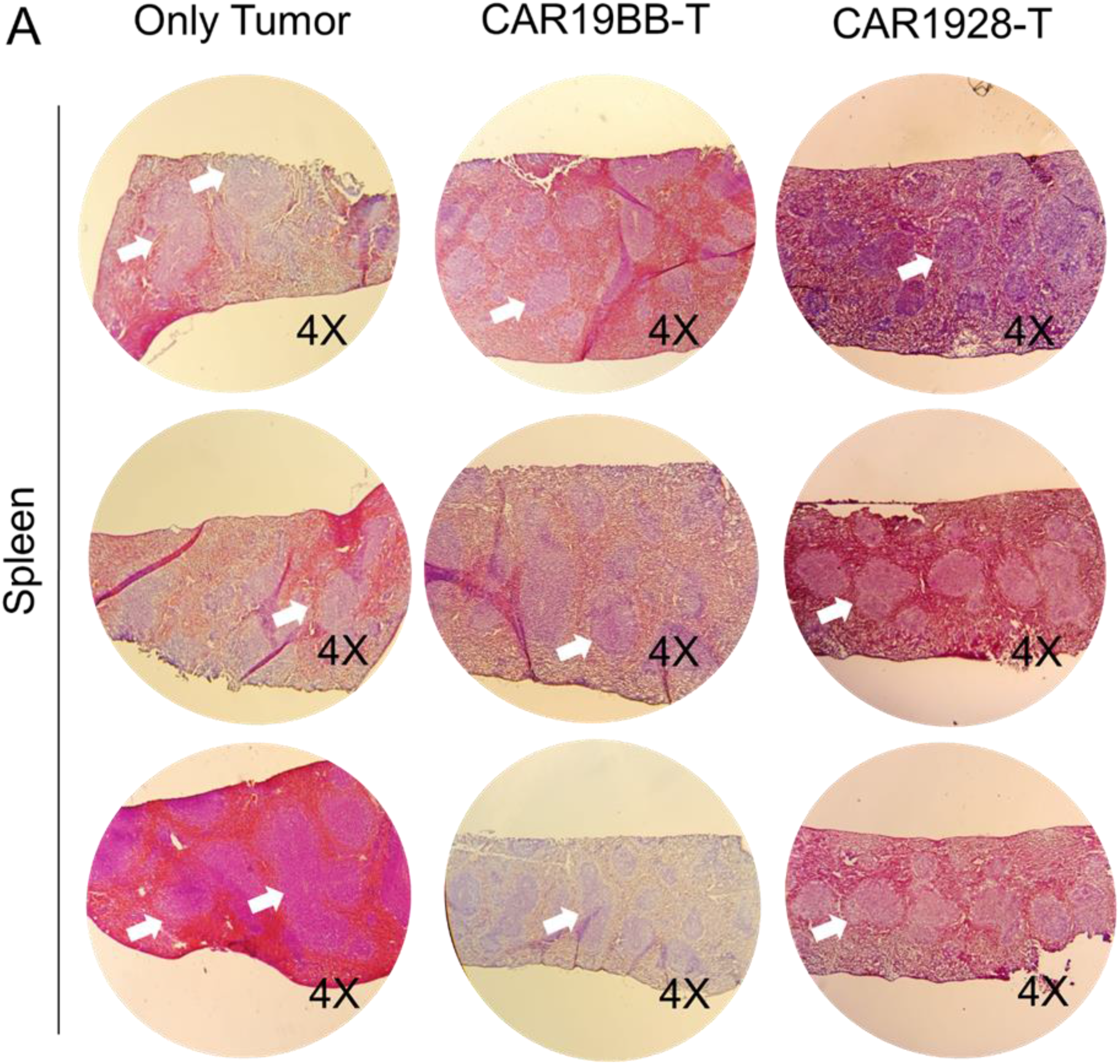
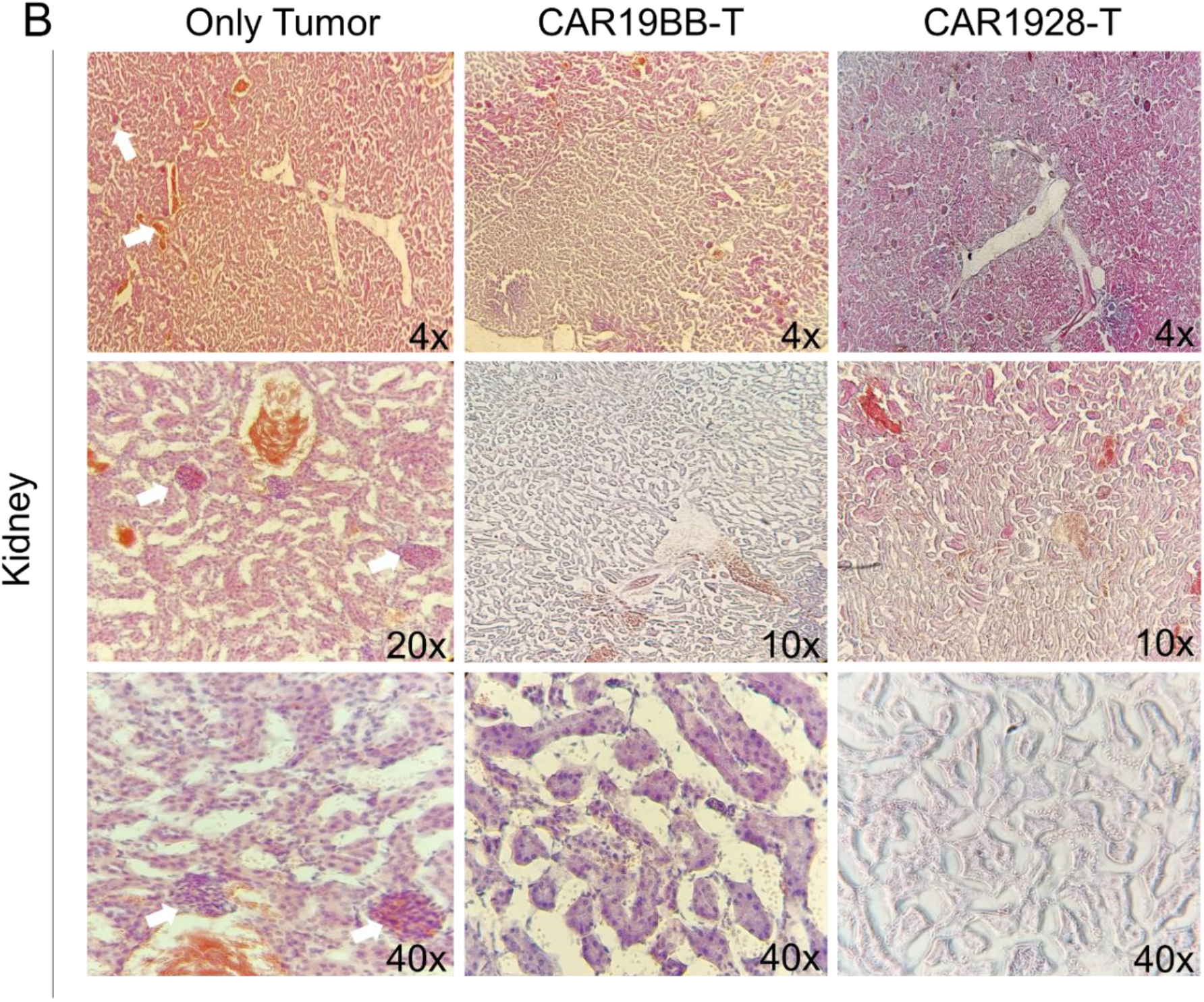
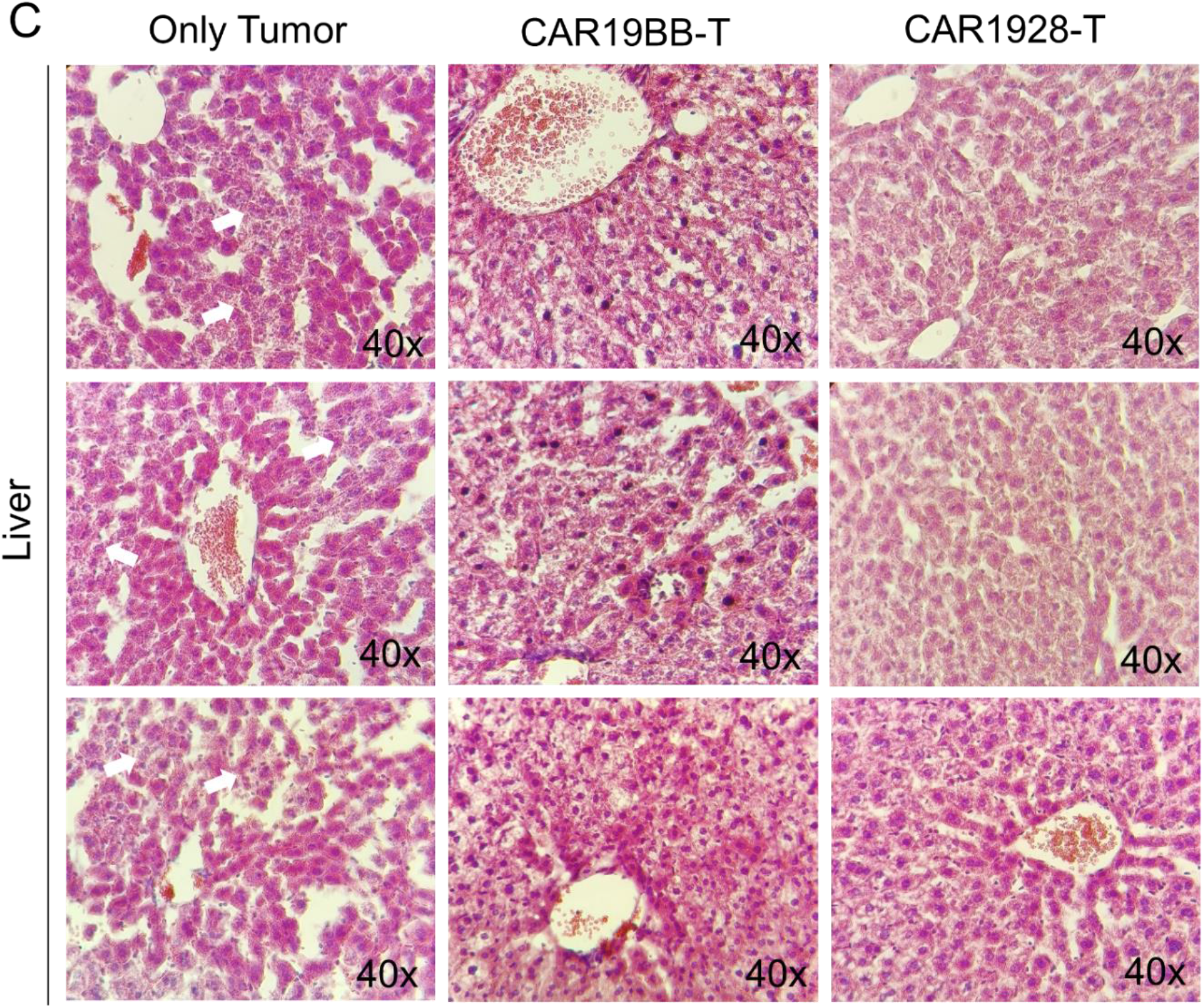
Histopathological analysis of the Spleen, Kidney, and Liver in the Only tumor and CAR-T cell groups. The images include inverted microscope views at 4X, 10X, 20X, or 40X magnification of (**A**) spleen, (**B**) kidney, and (**C**) liver sections obtained from the only tumor group sacrificed on day 35 and the CAR1928-T and CAR19BB-T-treated mice sacrificed on day 63 after Hematoxylin & Eosin staining.

## DISCUSSION

In some cancer treatments that require aggressive therapies, there is a need to improve the time-dependent efficacy and stability of CAR-T cells, which are a promising method. Current CAR-T cell technologies utilize anti-CD3/anti-CD28 microbeads to activate the cells. In this study, we report that PHA could be a viable alternative for activating and expanding CAR-T cells, resulting in improved stability and potential to transition into memory cell subtypes. Although current cellular immunotherapy treatments are ineffective in achieving complete remission, they do improve survival. One of the main reasons is that the patient’s immune cells have low-affinity receptors despite being stimulated against cancer antigens.

In this study, we investigated CAR-T production strategies using autologous T lymphocytes with high-affinity receptors and long-term stability. In order for CAR-T cells to remain long-lasting and effective in in vitro and in vivo cancer models, the proportions of TCM and TSCM sub-populations need to be high. We developed a new alternative CAR-T production process based on expansion using PHA instead of anti-CD3/anti-CD28 microbeads. Proliferation of PHA-activated T cells was significantly increased compared to activation with anti-CD3/anti-CD28 microbeads. Therefore, it appears that CAR-T cells activated with PHA can be expanded more rapidly and easily before being administered to patients in clinical trials. In both types of T cells, the TemEARLY cell population is significantly higher with PHA activation (Gulden et al., 2023). TemEARLY cells have a range of impressive functions and dominate in target tissues. They are more likely to reload tissue antigens compared to other T cell subpopulations. TemEARLY cells, due to their effector functions and long life, can contribute to the persistence of autoimmune diseases. The presence of a persistent antigen increases the proportion of TEM cell populations, as shown in studies on chronic infections; this also holds true for autoimmune diseases where self-antigens are continuously present (Devarajan, P. & Chen, 2013). The results of our in vitro and in vivo research suggest that TemEARLY cells generated after activation with PHA can persist in the bloodstream of patients as memory cells and provide a significant defense mechanism against recurring cancer (Gulden et al., 2023).

Our in vivo results can be interpreted to suggest that T cells expressing CAR1928 or CAR19BB, particularly those activated with PHA, demonstrate long-term anti-cancer capacity. In our animal studies using the ALL cancer model, CAR1928-T cells were effective in killing the injected CD19+ RAJI cells when used in conjunction with CAR19BB-T cells activated with PHA. In vivo analyses indicate that PHA may possess a similar or even superior capacity to anti-CD3/anti-CD28 in terms of effective anti-cancer capacity while providing higher long-term stability. Hemogram, biochemical, and histopathological analyses demonstrate that CAR19BB-T cells generated with PHA reduce necrosis and inflammation and improve tissue integrity in the mouse group, similar to CAR19BB-T or CAR1928-T cells generated with anti-CD3/anti-CD28 (Nayak, B. N., & Buttar, H., 2015; Cesta, M., 2006; Sato et al., 2012).

The production of CAR-T cells with PHA does not have a negative impact on cell viability compared to anti-CD3/anti-CD28 activation. Therefore, the use of PHA as an alternative to anti-CD3/anti-CD28 for CAR-T cell production may be a parallel research area for future clinical trials. Consequently, we conducted in vivo anti-cancer studies in an ALL animal model using CAR-T cells produced with PHA. In vivo studies demonstrated that CAR19BB-T cells produced with PHA maintained stability over time, and no relapse of cancer was detected in the CAR-T cell-treated group.

On the other hand, it was observed that CAR-T cell populations decreased and cancer relapsed in the groups treated with CAR19BB-T or CAR1928-T cells activated with anti-CD3/anti-CD28. Although these CAR-T cells began to proliferate with relapse, the CAR19BB-T cells activated with PHA did not show any decrease, suggesting a delay in cancer relapse. In future clinical trials, we plan to leverage both the CAR19BB-mediated memory T cell population and the high anti-cancer capacity of CAR1928-mediated T cells. In conclusion, CAR-T cell production with PHA exhibits a comparable or even better proliferation capacity than CAR-T cells activated with anti-CD3/anti-CD28.

